# Heterogeneous multimeric metabolite ion species observed in LC-MS based metabolomics data sets

**DOI:** 10.1101/2022.03.15.484295

**Authors:** Yasin El Abiead, Christoph Bueschl, Lisa Panzenboeck, Mingxun Wang, Maria Doppler, Bernhard Seidl, Jürgen Zanghellini, Pieter C. Dorrestein, Gunda Koellensperger

## Abstract

Covalent or non-covalent heterogeneous multimerization of molecules associated with extracts from biological samples analyzed via LC-MS is quite difficult to recognize/annotate and therefore the prevalence of multimerization remains largely unknown. In this study, we utilized 13C labeled and unlabeled *Pichia pastoris* extracts to recognize heterogeneous multimers. More specifically, between 0.8% and 1.5% of the biologically-derived features detected in our experiments were confirmed to be heteromers, about half of which we could successfully annotate with monomeric partners. Interestingly, we found specific chemical classes such as nucleotides to disproportionately contribute to heteroadducts. Furthermore, we compiled these compounds into the first MS/MS library that included data from heteromultimers to provide a starting point for other labs to improve the annotation of such ions in other metabolomics data sets. Then, the detected heteromers were also searched in publicly accessible LC-MS datasets available in Metabolights, Metabolomics WB, and GNPS/MassIVE to demonstrate that these newly annotated ions are also relevant to other public datasets. Furthermore, in additional datasets (*Triticum aestivum*, *Fusarium graminearum, and Trichoderma reesei*) our developed workflow also detected 0.5% to 4.9% of metabolite features to originate from heterodimers, demonstrating heteroadducts to be present in metabolomics studies at a low percentage.

## 1. Introduction

The ambition of metabolomics is to annotate and quantify every metabolite of the biological system under investigation for a comprehensive picture of all biological processes. However, currently, we are nowhere near this ambitious goal. While technological advancements on both the instrumental as well as the data processing side have greatly improved in recent years, researchers and analytical chemists are still faced with many challenges in their experiments. Especially, the annotation of observed features, including those originating from metabolite- and background compound-adducts, is challenging. This leads to a situation where a vast amount of features remains without annotation (i.e., the detectable portion of the dark matter in metabolomics [1]). However, the extent of these features arising from interactions between analytes remains largely understudied.

While a range of software solutions enabling the annotation of homo-multimeric adducts (e.g., [2M+H^+^]^+^) exist [2–4], these rely on predefined mass differences, peak shape correlation, and/or feature intensity correlation. Thereby more complex multimers, such as heteromeric adducts (e.g., [M_a_+M_b_+H^+^]^+^) or multimers forming before electrospray ionization (ESI) can not be annotated using the same strategies.

Indeed, only limited studies on the extent of heteromeric adduct formation exist. Specifically, Mahieu and colleagues developed and applied the mz.unity algorithm [5] to annotate ~6% of their credentialled features [6] in a dataset produced from *Escherichia coli* as multimeric adducts (not differentiating between homo and hetero adducts) [7]. Specifically, mz.unity computes possible multimeric relationships between observed mz values without considering elution profiles. While this allows for detecting heteromeric relationships it also leads to a significant risk for false positives. However, except for one case, a heteroadduct formed from glutamate and nicotinamide adenine dinucleotide, they did not focus on validating and characterizing these multimeric species in detail.

With regard to multimers formed before electrospray ionization (ESI) (which does not necessarily allow for annotation via the relationship of coeluting ions) due to chemical reactions that happen spontaneously after sampling (e.g., during sample preparation or chromatographic separation) these also hold the risk of misinterpretation. Indeed the occurrence of reactions between biomolecules is widely known [8,9] and even reactions between biomolecules and solvents have been studied [10].

Most of the presented works on analyte interactions are based on semi-targeted approaches and involve standards, which severely restricts their search space. This is likely because distinguishing signals arising from analyte interactions as compared to unique analytes is difficult even for skilled analytical chemists. Furthermore, the large amounts of data and compounds detected with modern LC-HRMS instrumentation make this an unmanageable task. Consequently, to the best of the authors’ knowledge, only a few non-targeted approaches screening for reactions between biomolecules have been conducted. As a result, the extent to which the analysis of metabolite extracts analyzed via LC-HRMS is affected by analyte interactions is largely unknown.

In this study, we attempt to inspect the extent of multimer formation between biological molecules present in LC-MS data obtained with reversed-phase chromatography (RPC) or hydrophilic interaction chromatography (HILIC). This includes analyte interactions taking place during ESI as well as in the samples. To achieve this, a novel workflow employing fully 13C labeled and unlabeled metabolomes was designed and applied to *Pichia pastoris* extracts. Briefly summarized, the two isotopically different yeast extracts were mixed. Any multimeric molecular species that spontaneously formed in this mixed sample or during ESI could therefore consist of labeled or unlabeled subunits (as well as combinations thereof) creating distinct isotopic patterns in LC-HRMS analysis. Such patterns are clearly indicative for different multimeric species (see Figure 1 and Figure 2). An in-depth evaluation of the detected heteroadducts suggested that heteroadduct formation is favored by specific chemical classes. Ultimately, we demonstrated that heteromers are not only present in our but also other datasets and deposited MS2 spectra acquired for our heteroadducts in the GNPS library initiating, to the best of the authors’ knowledge, the first MS2 library for heteroadducts.

**Figure 1.**
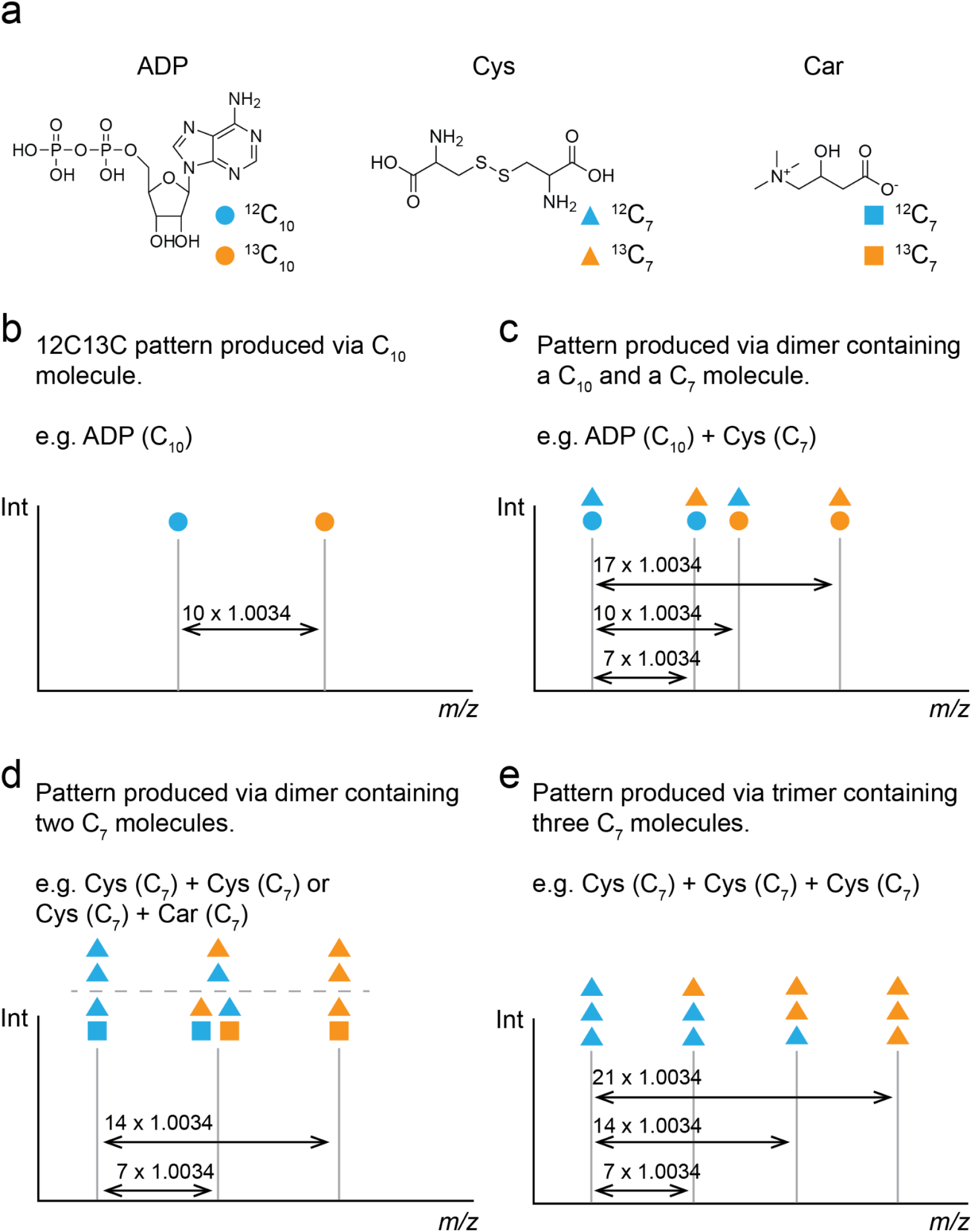
Examples for mz-patterns produced from mixtures of unlabeled and fully-13C labeled molecules. (a) Adenosine diphosphate (ADP), cystine (Cys), and carnitine (Car) present in their fully 13C labeled and unlabeled form within the same sample. (b) A molecule containing 10 carbon atoms. (c) mz-pattern of a heterodimer of a C10 and a C7 molecule. (d) mz-pattern of a dimer of two C7 molecules. (e) mz-pattern of a trimer of three C7 molecules

**Figure 2.**
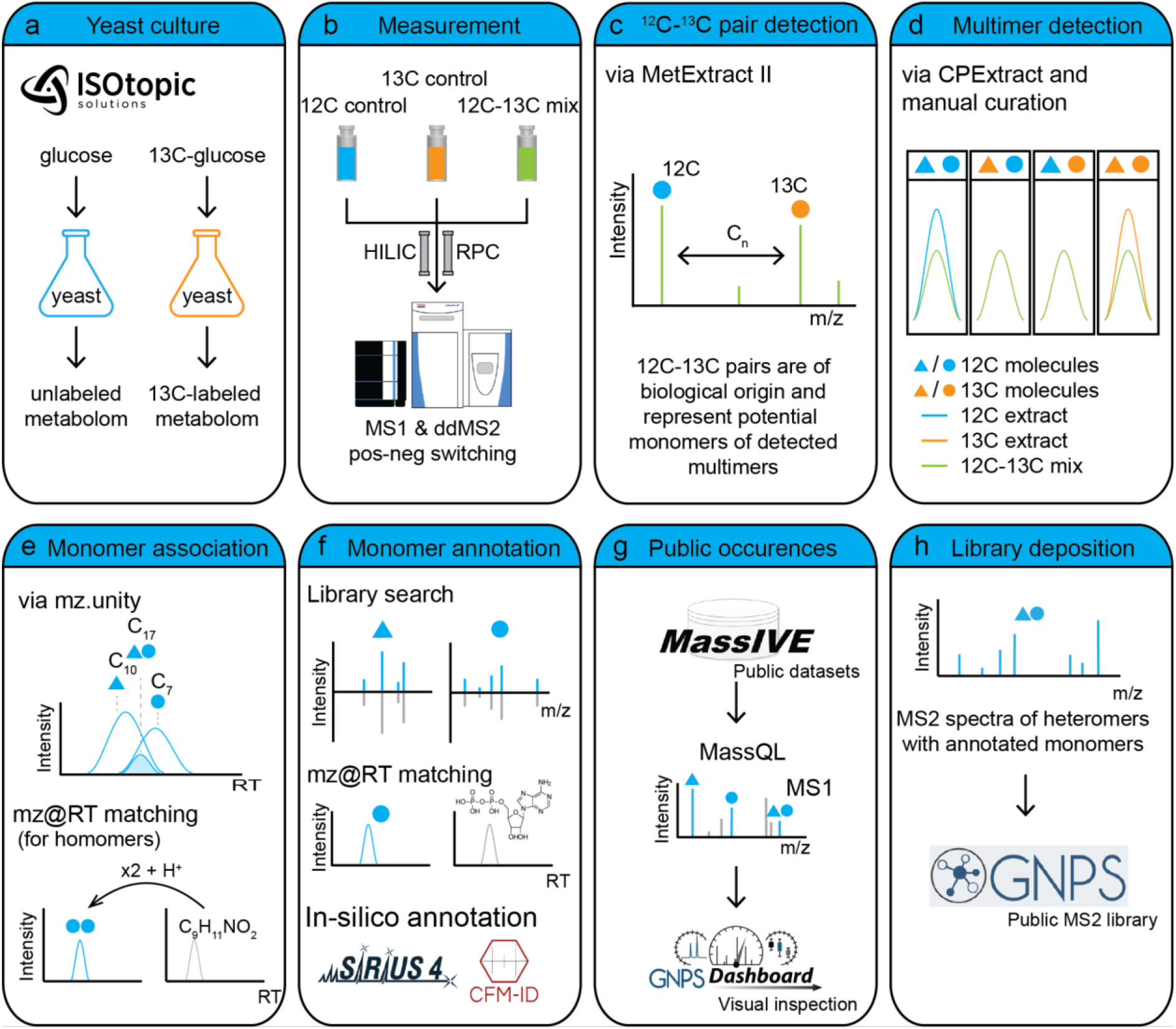
Individual steps of the workflow. (a) Yeast culture extracts were obtained from Isotopic Solutions. (b) 12C13C mixes and controls were analyzed via RPC and HILIC using MS1 and MS2. (c) Biological metabolites were detected as 12C13C pairs. (d) Multimers were detected via characteristic patterns within (see Figure 1) across samples. (e) Monomers were associated with multimers via mz.unity and RT matching to analytical standards. (f) Monomers were annotated via MS2 library searches, RT matching to analytical standards, and in-silico annotation. (g) Occurrences of multimers in public datasets were searched via MassQL and inspected via the GNPS dashboard. (h) MS2 spectra of detected multimers were deposited in the GNPS library for public use.

## 2. Materials and methods

### 2.1 Sample preparation

Unlabeled and uniformly 13C labeled ethanolic yeast (*Pichia pastoris*) extract pellets [11] were provided by Isotopic Solutions. Each was dissolved in 1 mL of LCMS grade water and centrifuged for 10 min. Three groups of samples were prepared from these two solutions. A 12C (only unlabeled yeast extract), 13C (only labeled yeast extract), 12C13C (mix of labeled and unlabeled yeast extracts), and a blank. Triplicates were prepared for each group. For RPC analysis, 60 μL of the 12C or the 13C samples were mixed with 140 μL LCMS grade water. For the 12C13C samples, 30 μL of the labeled extract, 30 μL of the unlabeled extract, and 140 μL of LCMS grade water were mixed. For HILIC analysis the same preparation was conducted but LCMS grade acetonitrile (ACN) instead of the LCMS grade water. For the blanks, LCMS grade water, and 70% ACN were used for both RPC and HILIC analysis, respectively.

### 2.2 Liquid chromatography

Samples were analyzed using a Dual-injection Vanquish liquid chromatography system (Thermo) coupled to a Q-Exactive-HF orbitrap mass spectrometer. As chromatographic columns, we used a Waters ACQUITY UPLC HSS T3 C18 column (2.1 x 150 mm, 1.8 μm) for RPC separation and a HILICON iHILIC-(P) Classic (2.1 x 100 mm, 5 μm) column for HILIC separation. In order to save time and eluents, and minimize instrumental variability between the RPC and HILIC separation we applied a previously published method to merge the RPC and HILIC separation into a single analytical run [12]. Using this setup we first eluted the HILIC and afterward the RPC into the MS. For RPC eluent A was 0.1% formic acid in LCMS grade water, and eluent B was ACN. For iHILIC eluent A was 15 mM ammonium acetate at a pH of 9.4 eluent B was ACN. For HILIC we injected 5 μL of sample and applied a flow rate of 0.2 mg mL^-1^. All gradient shifts were linear. The gradient was 90% B to 26% B (from 0 to 12 min), to 26% B (until 14 min), to 10% B (until 17 min), to 10% B (until 19 min), to 90% B (until 21 min), and at 90% B (until 36 min). At 15 min the valve was switched to the RPC column. Therefore, everything eluting past 15 min from the HILIC column was transferred into the waste instead of the MS. For the RPC we also injected 5 μL and applied a flow rate of 0.25 mg mL^-1^. During the first 15 min (while the HILIC analysis was still running) the RPC column was flushed into the waste at 0% B. At 15 min (when the sample was injected onto the RPC column) the amount of B was slowly increased to 99% B (until 29 min), stayed at 99% B (until 32 min), and was again decreased to 0% B (until 36 min).

### 2.3 Mass spectrometry

A Q-Exactive-HF orbitrap mass spectrometer was coupled to the liquid chromatography via an ESI source. All samples were analyzed in randomized order using MS1 and fast polarity switching for a mass range from 65 to 975 mz. The AGC target was set to 1e6 counts, the maximum injection time to 100 ms, and the resolution to 60000 at 200 *m/z*. Afterward, the 12C13C-pair and multimer pattern extraction was conducted as described below. The two obtained feature lists were combined and their mz and retention time values served as targets for inclusion lists for subsequent ddMS2 acquisition. MS2 spectra were acquired at a resolution of 30000, stepped collision energy (30, 50, 80 NCE), and with an isolation window of 1 mz for positive and negative polarity.

### 2.4 12C13C-pair extraction

Biological metabolite credentialing was carried out with MetExtract II [13]. The software automatically searched LC-HRMS datasets, converted to centroided mzXML format via ProteoWizard [14], for pairs of unlabeled and uniformly 13C labeled metabolite ions and verified them on both the mz and retention time dimensions. In the mz dimension, it checked if both isotopologue forms showed their characteristic isotopologue patterns that originate from the 98.93% 12C and 1.07% 13C isotope composition in the unlabeled samples and from the 98.7% 13C and 1.3% 12C isotope composition in the uniformly 13C labeled samples as determined automatically from the isotopologue patterns for some 2100 features. Moreover, as unlabeled and 13C labeled isotopologues showed no chromatographic separation, high similarity in the chromatographic peak shapes of the principal isotopologues was tested for as well. Furthermore, after detection of such pairs of unlabeled and 13C labeled metabolite ions in the different samples, MetExtract II grouped/aligned these features across the different samples, convoluted these different features into groups each representing a certain truly biologically-derived metabolite, and re-integrated the principal isotopologues in the 12C, 13C, or blank samples to generate a comprehensive data matrix. MetExtract II reported the mean intensity in the chromatographic peaks’ apex (i.e., the mean intensity value of the peak apex and its two preceding and succeeding scans). Specific parameter settings were: Cn count to search for: 3 – 60, retention time domain to search for: 15 – 40 minutes for RPC method and 1 – 15 minutes for HILIC, minimum intensity threshold for signals: 1E4 (only applicable for the monoisotopic and the fully 13C labeled principal isotopologues), maximum allowed mass deviation for isotopologues in the same scan: 6 ppm, isotopologue peaks checked: M + 1 and M + Cn-1, maximum cluster ppm for the region of interest (ROI) generation: 8 ppm, minimum number of signals with verified isotopologue pattern: 3, chromatographic peak width in wavelet terms: 1 – 3, bracketing mz and retention-time window: 12 ppm and 0.1 minutes, minimum Pearson correlation coefficient (PCC) for feature grouping were 0.85.

### 2.5 Multimer pattern extraction and curation

CPExtract is a software tool for the processing of LC-HRMS datasets derived from stable isotope labeling experiments [15]. It allows the user to freely define custom isotopologue patterns in the form of presence/absence and intensity rules. Each signal recorded in the LC-HRMS data is tested with these rules and those fulfilling them are further considered. Thereby, all isotopologue patterns defined by the user are detected and any isotopologue patterns that are not within the defined boundaries are discarded. Moreover, as a second step CPExtract also performs chromatographic peak picking, grouping/alignment of the features detected in the different samples, convolution of features of the same metabolite as well as re-integration of features in blank samples.

To search for homo- and hetero-multimers two different rule sets were used:

Homomers: For homomers consisting of two molecules of the same compound, the rules were set up in a way that three isotopologue patterns were checked. A variable number of carbon atoms n was allowed. The principal isotopologues were required to have the mz values X, X+ΔC*n, and X+ΔC*2n, with ΔC being 1.00335 (i.e. the mass difference of 13C and 12C). Moreover, the absence of signals at M-ΔC and M+ΔC*(2n+1) was assured. Additionally, M+ΔC had to be less abundant than M, M+ΔC*(n-1) and M+ΔC*(n+1) less abundant than M+ΔC*n, and M+ΔC*(2n-1) less abundant than M+ΔC*2n. Heteromers: For heteromers, the rules were set up in a way that four isotopologue patterns were demanded. A variable but an unequal number of carbon atoms a and b were allowed. The principal isotopologues were required to have the mz values X, X+ΔC*a, X+ΔC*b, and X+ΔC*(a+b). Similar to the homomers, also no signals were allowed at X-ΔC* and X+ΔC*(a+b+1). Furthermore, also the isotopologues surrounding the principal isotopologues must be less abundant than the respective principal isotopologues.

Additional parameters other than these two separate rule sets were identical with the parameters of the MetExtrat II data evaluation for metabolite credentialing.

After multimer detection via CPextract, the absence of isotopologue chimers in 12C and 13C controls, and all multimer-isotopologues in solvent-blanks was confirmed via R scripting.

### 2.6 Monomer association

The association of multimer patterns to their respective monomers was conducted by two different methods. First, for homomers, an in-house retention time (RT) library including molecular formulas with known RTs was applied. Using a list of possible multimeric adducts a range of mz values were calculated and matched to the observed multimers with an RT tolerance of 0.2 min and a mass tolerance of 5 ppm. The comprehensive list of adducts is given in Table S1. For heteromers and homomers for which this strategy did not succeed we applied mz.unity [5]. In mz.unity all 12C13C-pairs eluting within 0.2 min of a given multimer were considered as monomer candidates. Considered charge carriers are given in Table S2. For all association methods candidates were only accepted if the carbon number of the 12C13C-pairs fit the number of carbons required by the respective multimeric adducts. Finally, in cases where the PCC-grouping of hetero-multimers was in conflict with monomer association, PCC groups were split.

### 2.7 Annotation of monomers and multimers

Putative monomers were annotated using a range of different approaches. First, in cases where the monomer association was achieved by retention time library matching (as described above), the annotation was already implied by the monomer association. However, we also applied our RT library for the identification of monomers in the same manner as described in section 2.6. Moreover, MS2 library matching of acquired MS2 scans was done using two different fragmentation libraries. For the GNPS library, we used the GNPS infrastructure for spectral matching [16]. The precursor ion mass tolerance was set to 0.025 Da, min matched peaks to 3, fragment ion mass tolerance to 0.05 Da, and the score threshold to 0.7. For the mzCloud library matching Thermo Scientific’s Compound Discoverer 3.1 was applied. The precursor and fragment mass tolerance were set to 10 ppm, and the match factor threshold to For heteromer monomers that remained unannotated, but an MS2 spectrum was acquired, Sirius [17] or CFM-ID 4.0 [18] were applied. For Sirius and CSI::FingerID (within Sirius), the instrument type was set to “Orbitrap” and all other values to default. In cases where Sirius was not able to deliver a result, CFM-ID 4.0 was applied. The candidate mass tolerance was set to 10 ppm, and all other values were at default. Finally, for all annotation methods described above candidates were only accepted if their carbon counts fit the number of carbons determined during 12C13C-pair (or multimer pattern) extraction.

### 2.8 Other inspected datasets

Details for the analysis and data evaluation of the other inspected datasets (*Triticum aestivum*, *Fusarium graminearum, and Trichoderma reesei*), are given in the Supplementary information (SI 1-4) [10,19,20].

### 2.9 MassIVE database inspection

The MassIVE database was inspected via tools of the GNPS infrastructure MassQL (for MS1 based patterns) and MASST [21] (for MS2 spectral matching). Regarding MassQL we required MS1 scans in the repository to include the mz values of the heteromer as well as the two monomers associated in our study with a tolerance of 20 ppm. Subsequently, we applied R to remove MassQL hits if they did not occur in at least contiguous 4 scans of the same polarity. All remaining hits were manually inspected and only accepted if the heteromer showed a high correlation with at least one of the monomers. For the MASST search, we applied a parent mass tolerance of 0.05 Da, an ion tolerance of 0.05 Da, and a minimum of 3 matching fragments.

## 3. Results and Discussion

### 3.1 The developed workflow

Multimer formations can occur during sample preparation or ESI. Such products and multimeric adducts typically have strikingly similar characteristics to any other compound also recorded for the sample (i.e., a chromatographic peak and corresponding *m/z* distribution). This bears the risk of mistaking them for individual biological molecules.

The workflow for the detection of chemical reaction products in untargeted LC-HRMS-based approaches is based on the utilization of ethanolic extracts from uniformly 13C labeled organisms and their respective unlabeled counterparts (Figure 2 a). Samples of both, the 13C labeled and native organisms, are mixed into a single sample, which allows to easily discriminate between metabolites and background compounds, a process often referred to as credentialing. Any metabolite will be present as an unlabeled and a 13C labeled form likewise, but not as any mixed form since the isotopic markup of the respective molecules has been established during the separate cultivation of the 13C labeled and unlabeled organisms (Figure 1 a, b).

Contrary, metabolite-multimers formed after mixing the labeled and unlabeled extracts will show not only these two distinct isotopic patterns (the one originating from the unlabeled and another from the 13C labeled cultivation) but additional ones that are indicative for multimers. Multimers of two identical molecules or two different molecules with the same number of 13C labeled carbon atoms will show three characteristic isotopic traces, while multimers of two different metabolites with different numbers of carbon atoms will show four characteristic isotopic traces (Figure 1 c and d). Thus, automated data analysis of the LC-HRMS dataset will allow finding such multimers even in an untargeted fashion.

To demonstrate this approach, extracts of yeast grown separately under unlabeled and uniformly 13C-labeled conditions were employed. Their extracts were mixed and these mixtures were analyzed with reversed-phase chromatography (RPC) and hydrophilic interaction chromatography (HILIC) coupled to high-resolution mass spectrometry (Figure 2 b). Multimers of molecules that were present in their labeled and unlabeled form were then detectable via specific isotopologue patterns, as exemplified in Figure 1. The detection and validation of these patterns were performed via a combination of the CPExtract software and manual curation (Figure 2 c). To associate the detected multimers with their monomers, we used the MetExtract II software to generate a list of 12C-13C feature pairs (Figure 2 d), representing yeast-derived metabolites, which served as a list of possible monomer candidates for the detected multimers. The association of these candidates with the multimers detected was then conducted via the mz.unity package (Figure 2 e). This approach was complemented with our in-house library of analytical standard compounds, which allowed us to match mz@RT-features annotated as homomultimers directly against a list of putative homomultimers (e.g., [2M+H^+^]^+^, [3M+H^+^]^+^, [2M+Na^+^]^+^, etc.) (see Table S1 for a comprehensive list of the applied search space). Since this way of association did not require the coelution of monomers it also enabled the annotation of multimers formed during sample preparation instead of during ESI.

The annotation of putative monomers was conducted by matching associated MS2 scans to the GNPS and mzCloud mass spectral libraries. Further, we applied our in-house RT library as described in section 2.6, and in-silico tools, if the first two approaches did not succeed (Figure 2 f). To assess whether observed heterodimers were also detected in other labs and studies, we used MassQL, which allowed querying the MassIVE database (also including datasets deposited on Metabolights [22] and Metabolomics WB [23]) for scans where the mz-values of a heteromer and its associated monomers appeared in the same MS1 scan (Figure 2 g). Several hits have been manually confirmed and can be inspected via the GNPS dashboard [24]. Finally, MS2 scans of identified multimeric adducts were deposited in a GNPS MS2 library so that their fragmentation scans can be automatically annotated in future studies (Figure 2 h).

### 3.2 Detection of biological signals in yeast data

MetExtract II allows the automatic extraction of 12C-13C-pairs (mz@RT values of compounds that are present in their unlabeled and fully labeled form). In our case, these 12C-13C pairs corresponded to biological metabolites truly derived from yeast as contaminations are only present in their unlabeled form but not as fully 13C labeled isotopologues. In this study, we detected 923 and 627 peaks representing metabolite features with known carbon counts that were convoluted automatically with 332 and 288 metabolites for HILIC and RPC, respectively. These features served as possible monomers since the detected multimer pattern required the associated monomers to be present as labeled and unlabeled forms.

### 3.3 The frequency of multimeric LC-MS peaks

CPExtract extracted isotopic patterns specific for metabolite-multimers, formed after mixing of the labeled and unlabeled extracts, irrespective of their origin (e.g., ESI or sample preparation). As visualized in Figure 1 (c, d), two different types of patterns were searched for. Those patterns were indicative of the presence of a dimer composed of two carbon-bearing molecules with the same or a different number of carbon atoms, respectively. Using this approach, we were able to detect 33 homodimers and 7 heterodimers in the RPC and 68 homodimers and 14 heterodimers in the HILIC analysis of the yeast samples. All detected patterns were subsequently manually inspected and duplicate detections of the same patterns and other data processing artifacts were removed. Moreover, 3 patterns associated with heteromers (Figure 1 c) were manually reassigned to homomultimers consisting of more than two monomers (e.g., trimers and tetramers), which were incorrectly assigned via CPExtract (see Figure 1 e). The application of monomer association strategies revealed that 3 of the homomers detected for HILIC were actually heterodimers of monomers with the same numbers of carbons. Consequently, the number of detected patterns changed to 26 homodimers and 5 heterodimers for RPC and 50 homodimers, 14 heterodimers, and 3 homomultimers for HILIC, respectively.

In summary, 5.7% of peaks of biological origin (53 peaks / 923 12C-13C pairs) could be attributed to homomers when HILIC separation was employed. For the analysis employing RPC, homomers were slightly less prevalent at 4.1% (26 peaks / 627 12C-13C pairs). Interestingly, for HILIC only 1.5% of peaks with biological origin (14 peaks / 923 12C-13C pairs) could be attributed to heteromers in our experiment. For RPC this percentage was even lower with 0.8% (5 peaks / 627 12C-13C pairs). The low percentage of heteromeric peaks detected holds important implications as currently no software solution for their routine annotation in LC-MS experiments exists. The complete lists of detected homo and heteromers are provided in Table S3 and Table S4.

To elucidate the number of biological molecules contributing to the detected multimers, we investigated the peak shape correlation between the detected multimeric species. If the Pearson Correlation Coefficient (PCC) between a set of multimers was above a threshold they were considered to originate from the same molecule but as a different ion species (e.g., different charge carriers) and assigned into the same PCC-group. This resulted in 32 and 17 biological molecules contributing to homomeric species for HILIC and RPC, respectively. For heteromeric species we conducted the same grouping, leading to 12 (HILIC) and 4 (RPC) PCC groups. Notably, in the case of heteromers, different ion species could be a consequence of one of the monomers being substituted by an alternative monomer.

In general, we focus on heteromers throughout most of the discussion, as homomers are readily annotatable by many non-targeted metabolomics tools (e.g., ion identity networking, or CAMERA). For heteromers, on the other hand, no readily applicable solution exists, which leads to a situation where the risk of their misannotation as unique metabolites is high.

### 3.4 The origin of the detected multimers

Multimers can form at different stages of the analytical workflow (e.g., during the sample preparation, storage, chromatographic separation, or ESI). To elucidate if multimers formed during the ESI, we assessed whether the detected multimers could be explained via co-eluting metabolites. Overall, we were able to associate co-eluting monomers for 57% of heteromers (8 of 14) and 49% of homomers (26 of 53) for HILIC and 60% of heteromers (3 of 5), and 84% of homomers (21 of 26) for RPC to monomers.

Another way to investigate the origin of homomeric dimers is to scrutinize their intensity patterns. Any homodimer of biological origin (i.e., formed or bio-synthesized by the organism) can only be present as a purely non-labeled or a purely isotopically labeled form (Figure 1 b). Thus, intermediate patterns representing mixed forms (i.e., chimeric isotopologues with one of the educts being unlabeled while the other one being labeled) will not be present. On the other hand, any dimer that forms exclusively after the unlabeled and labeled extracts are mixed will show a distinctive isotopologue distribution that depends on the abundance of its educts (see Figure S1 a). Thus the comparatively low abundance of chimeric isotopologues allows inferring the contribution of biological as well as post-extraction contribution to a multimeric species (i.e., Figure S1 b), which excludes the ESI-process as multimer-origin. In total, we found that 17% (9/53) and 4% (1/26) of the discovered homodimers fell within this category for HILIC and RPC, respectively. The fact that these numbers were different for the different separations implies that the formation of these homodimers was impacted by the sample solvents (70% acetonitrile (HILIC) or water (RPC)) or chromatographic conditions (e.g., pH = 9.2 (HILIC) or pH ≈ 2 (RPC)). It should be noted that irreversible reactions which reached equilibrium before the labeled and unlabeled extracts were mixed were not picked up by this strategy.

Applying both described strategies and PCC-grouping we were able to elucidate the origin of 70% and 92% of multimer origins for HILIC and RPC, respectively. Multimers that could not be associated with co-eluting monomers could be explained by unexpected charge carriers (for ESI), bioinformatic errors (such as those caused by potential problems in feature finding), or fast solution reactions.

### 3.5 Identification of monomers

In the following, we describe the results of our attempts to identify the monomers, which have been associated with the detected PCC-groups for ESI-adducts and reaction products. Identification was conducted via retention time matching to our in-house standard library, MS2 matching to fragmentation libraries (GNPS and mzCloud), and annotation via in-silico tools (CSI::FingerID and CFM-ID). In Figure 3 we summarized which molecules have been annotated thereof. Overall, we were able to annotate 100% (12/12) and 83% (10/12) of monomers associated with homomer PCC-groups for HILIC and RPC, respectively. Regarding heteromers, we annotated 100% of the associated monomers (14 for HILIC and 6 for RPC).

**Figure 3.**
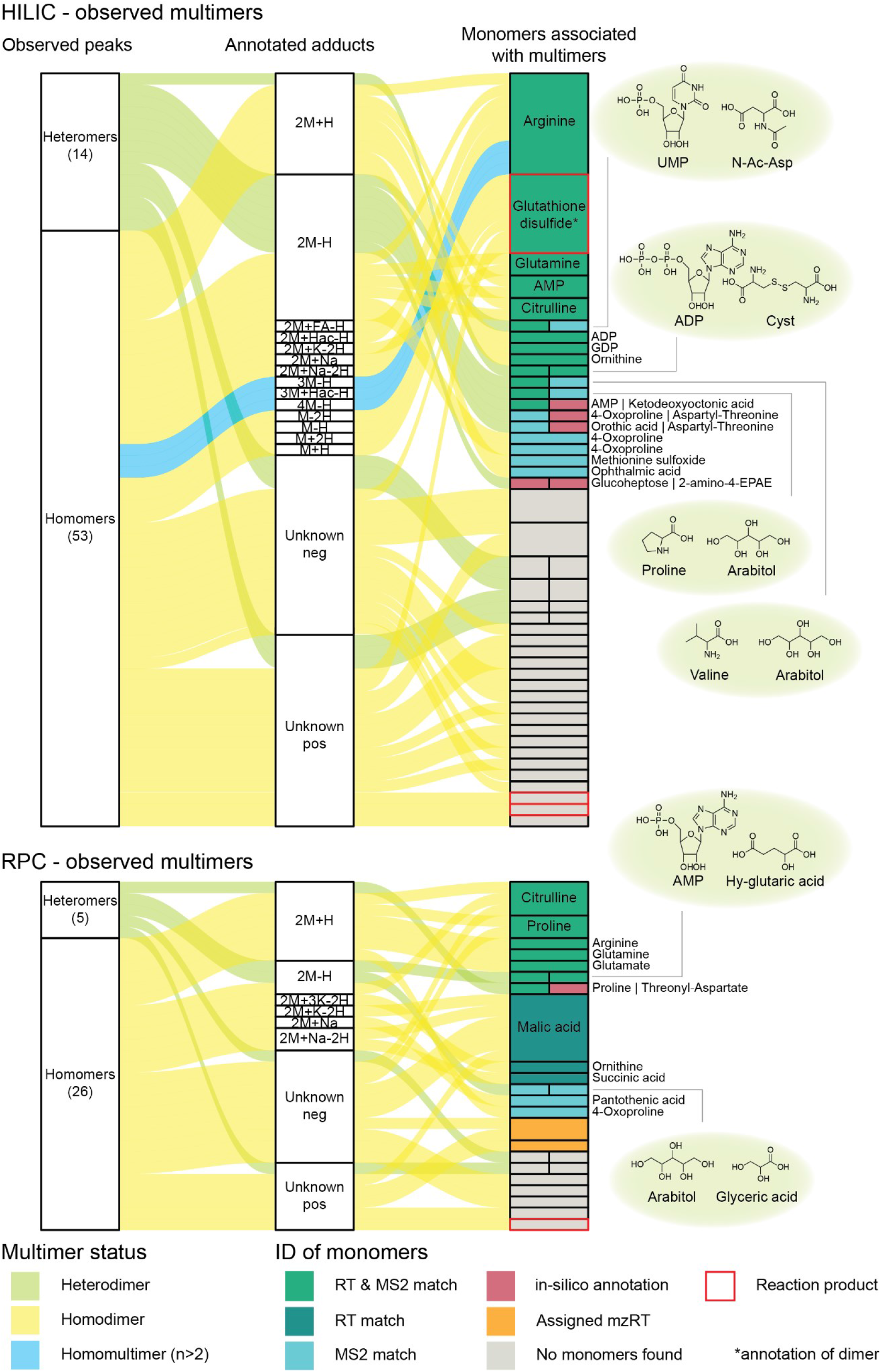
plot of monomer-multimer relationships. (left box) Hetero- and homomers detected after RPC and HILIC analysis. (middle box) Observed multimeric adducts. (right box) Associated PCC-groups with monomer annotation and annotation source.

Notably, 4 of 11 heteromers for which monomers were identified included a nucleotide (AMP, UMP, or ADP), and 3 of 11 Arabitol. This enrichment of specific metabolite classes in the formation of heteromers suggests that the classes of compounds present in a given analyzed sample (and co-ionizing during ESI) have a significant impact on the frequency of observed heteroadducts.

Another interesting aspect of the conducted study was that only 3 multimers consisting of more than 2 monomers were detected in the HILIC dataset (as noted in section 3.3). All three of these multimers turned out to originate from Arginine and were detected as [3M-H^+^]^−^, [3M+acetic acid-H^+^]^−^, and [4M-H^+^]^−^ adducts, suggesting that multimeric adducts (n>2) might indeed be quite rare.

### 3.6 Relevance of findings for the metabolomics community

#### 3.6.1 Properties of detected multimers

The intensity of observed multimers is an important metric for judging their impact on metabolomics results. As can be seen from the histograms illustrated in Figure 4, the majority of observed multimers (and the vast majority of heteromers) were less abundant than the median of biological peak abundances. This reduces the likelihood of these ions triggering fragmentation scans in a classical Top-N ddMS2 experiment in which the N most abundant ions of a given MS1 scan serve as the precursors for subsequent MS2 scans. Consequently, these multimers are unlikely to impact many non-targeted metabolomics strategies such as molecular networking and other MS2 scan-based strategies, if not specifically targeted during MS2 scan acquisition. Nevertheless, their occurrence bears the risk of misinterpreting them as unique biological metabolites on the MS1 level. Moreover, newer methods and instruments keep increasing the MS2 coverage achievable within metabolomics experiments [25–27] which increases the chances of acquiring MS2 scans also for these low abundant ion species.

**Figure 4.**
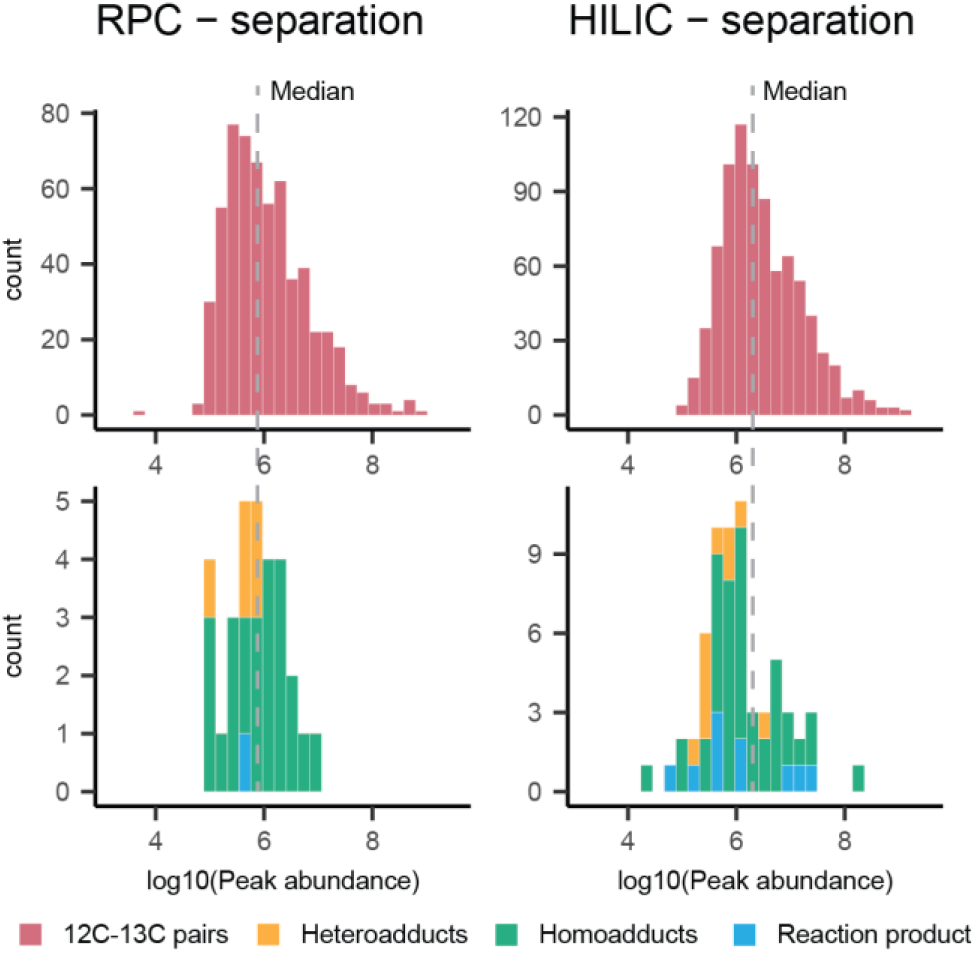
Histograms of peak abundance distributions. Peak abundance distributions for metabolites (12C13C pairs) and multimers colored by their origin for RPC and HILIC. The dashed line indicates the medians for metabolites of both separations.

The PCC correlation between multimers and respective simpler adducts (such as [M+H^+^]^+^ or [M-H^+^]^−^) is of importance for their grouping in classical metabolomics workflows (as possible via software such as CAMERA or Ion Identity Networking). Figure 5 gives an overview of this important metric for multimeric adducts. Figure 5 a shows that the majority of the detected homomers show good peak shape correlation with the respective [M+H+]^+^ or [M-H+]^−^ adduct for HILIC and RPC. However, in the case of heteromers, only one monomer shows a good peak shape correlation with the respective heteromer in most cases (see Figure 5 b). This emphasizes that relying on the PCC alone is suboptimal for grouping heteroadducts with their respective monomers. However, this finding provides a potential starting point for the association of heteroadducts with their respective monomers in classical non-targeted experiments where stable isotope-labeled material is not available. Extracted ion chromatograms of all annotated heteromers and their associated monomers are provided in Figure 6 for closer inspection.

**Figure 5.**
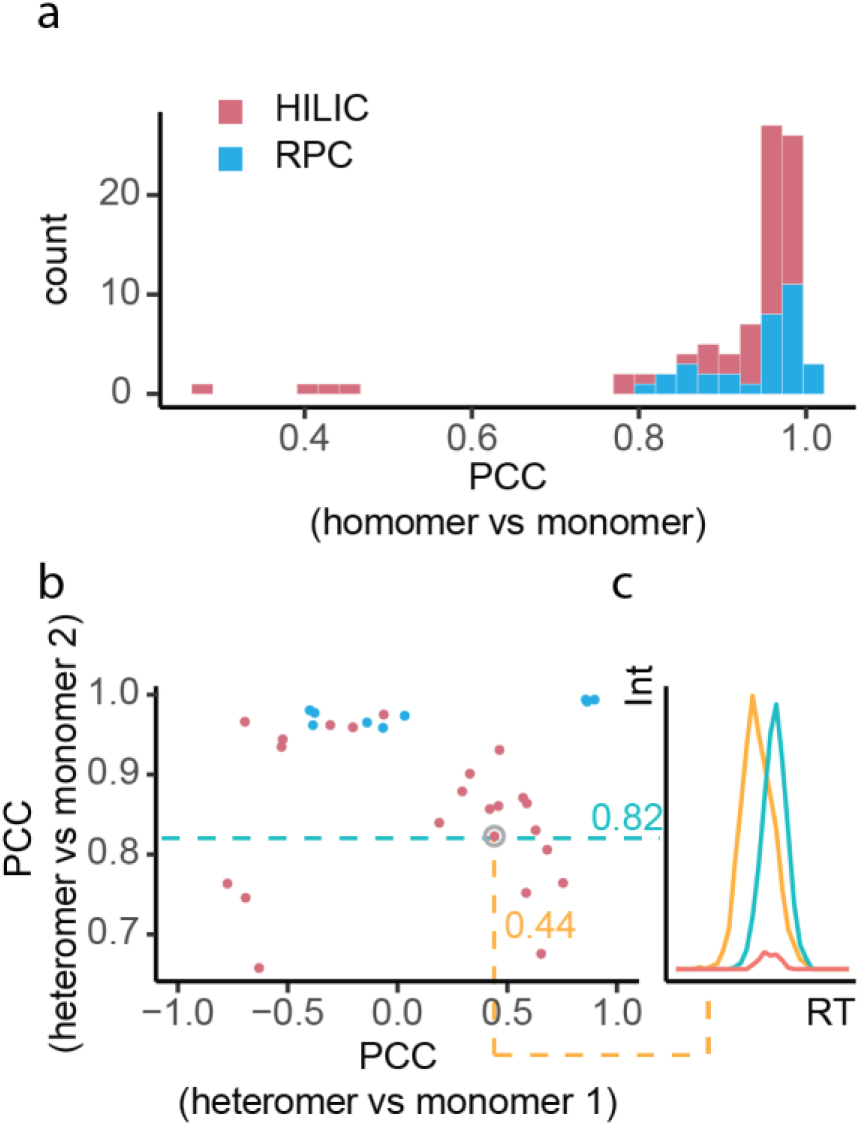
Distribution of Pearson Correlation Coefficients (PCC) of multimers with their monomers. (a) PCC of homomers with associated monomers. (b) PCC of heteromers with both monomers. (c) Example for heteromer elution profile (red) as compared to monomer elution profiles (orange and blue).

**Figure 6.**
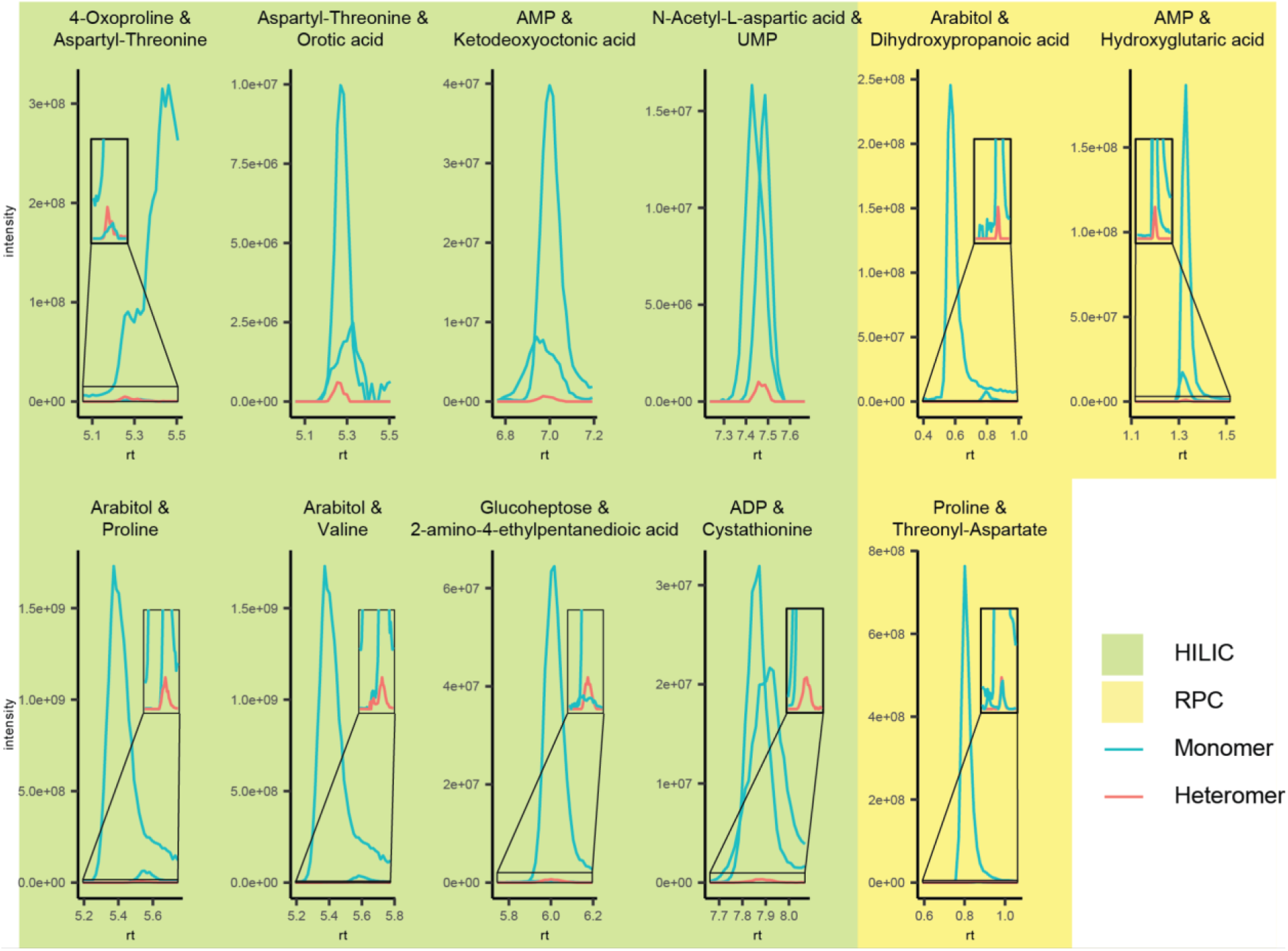
Extracted ion chromatograms for all annotated heteromers and their associated monomers. For RPC retention times the time of the sample injection on the chromatographic system was set to 0.

#### 3.6.2 Occurrence of observed heteromers in other data sets

In order to estimate whether our findings hold implications for other LC-MS datasets implicating other sample types and instrumental platforms, we reanalyzed three datasets involving different fully 13C labeled organisms (*triticum aestivum, fusarium graminearum,* and *Trichoderma reesei*) on a different instrumental platform. Specifically, percentages of multimeric species in these other datasets were assessed in the same way as the yeast dataset. As can be seen in Table 1, although commonly observed, the percentages of heteromeric features did not exceed 4.9% of all biological features in any of the presented data sets. This suggests that most of the complexity obtained in LC-MS mass spectra is usually not caused by excessive multimeric adduct formation of biological analytes.

**Table 1.** Detected features and PCC-groups.

| Dataset - Separation principle | Biological metabolites<sup>1</sup> | Homomers<sup>2</sup> | Heteromers<sup>2</sup> |
| --- | --- | --- | --- |
| *Pichia pastoris* - HILIC | 923 / 332 | 54 / 34,<br>5.9% / 10.2% | 14 / 12,<br>1.5% / 1.3% |
| *Pichia pastoris* - RPC | 627 / 288 | 26 / 17,<br>4.2% / 5.9% | 5 / 5,<br>0.8% / 0.8% |
| *Triticum aestivum* - RPC | 2354 / 590 | 170 / 97,<br>7.2% / 16.4% | 116 / 90,<br>4.9% / 15.2% |
| *Fusarium graminearum* - RPC | 1519 / 554 | 24 / 17,<br>1.6% / 3.1% | 8 / 8,<br>0.5% / 1.4% |
| *Trichoderma reesei* - RPC | 1856 / 779 | 45 / 25,<br>2.4% / 3.2% | 17 / 15,<br>0.9% / 1.9% |

Number of biological features and metabolites and multimers detected in different datasets. 1... numbers stated as: number of metabolite features / number of metabolites, 2... numbers stated as: number of features / number of PCC-groups, percent relative to the total number of metabolite features / percent relative to the total number of metabolites

Finally, we assessed whether heteroadducts identified in the presented yeast data set have been observed in publicly available LC-MS data deposited at the MassIVE repository. Using MassQL we assessed whether mz values of the annotated heteroadducts appeared within the same MS1 scan as their associated monomers (as listed in Table S4) in at least 4 consecutive MS1 scans of the same polarity. After manual curation, we concluded that there was sufficient evidence suggesting that 4 of the heteromers identified in our data set were also present in data sets produced by other labs analyzing samples originating from organisms such as *escherichia coli*, marine plankton, *caenorhabditis elegans*, and *rattus norvegicus*. A full table including all heteromers detected in MassIVE as well as links to interactive extracted ion chromatograms (as facilitated by GNPS Dashboard) of these hits is given in Table 2. Notably 2 of the 4 public datasets in which heteromers were discovered applied HILIC. This is notable as according to REDU [28] less than 2% of MassIVE datasets applied HILIC, while most used RPC. As we found significantly more multimeric species on HILIC as compared to RPC in our yeast datasets, this further supports the hypothesis that HILIC seems to be more prone to heteromultimer formation.

**Table 2.**
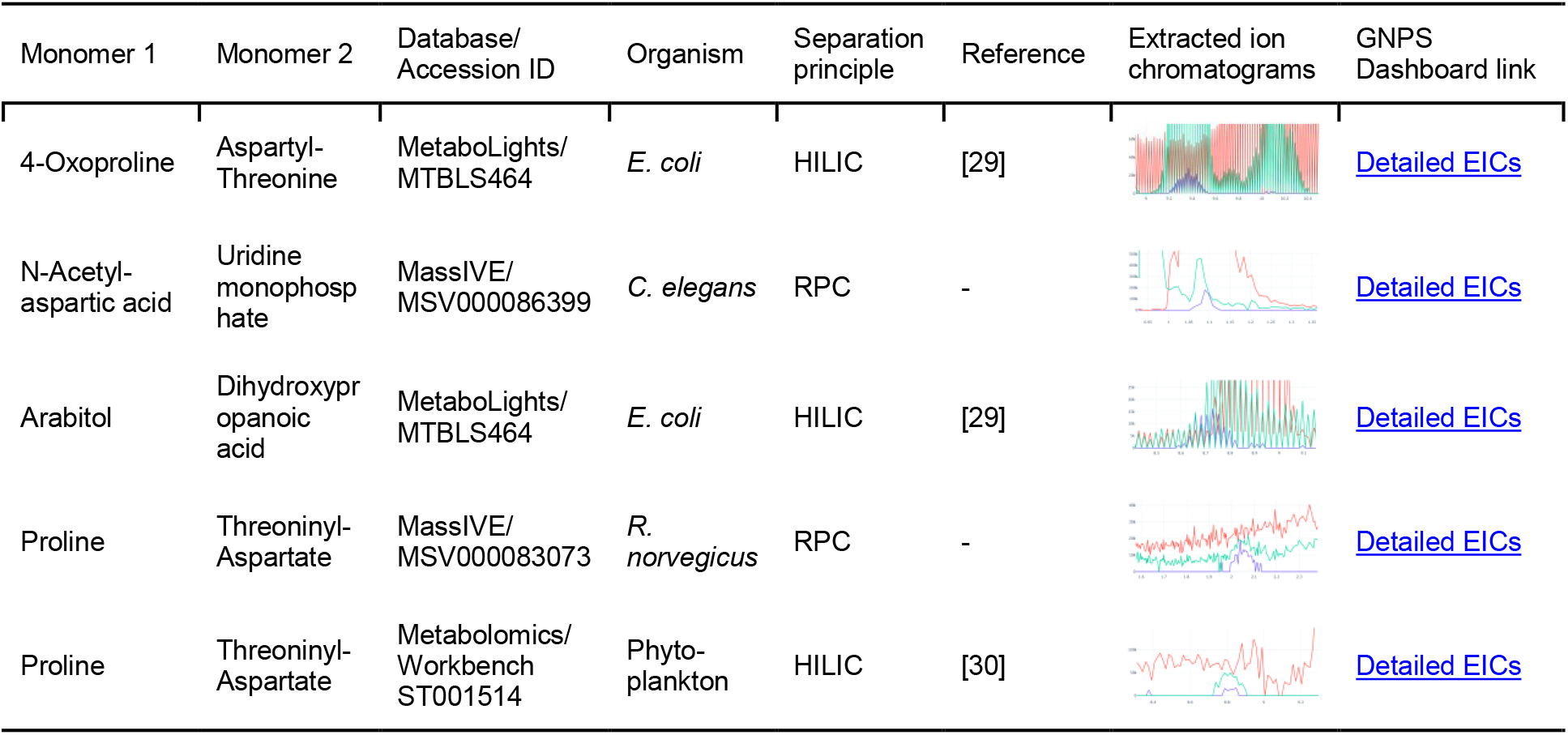
Heterodimers found in public datasets.

| Monomer 1 | Monomer 2 | Database/<br>Accession ID | Organism | Separation<br>principle | Reference | Extracted ion<br>chromatograms | GNPS<br>Dashboard link |
| --- | --- | --- | --- | --- | --- | --- | --- |
| 4-Oxoproline | Aspartyl-Threonine | MetaboLights/<br>MTBLS464 | *E. coli* | HILIC | [29] |  | Detailed EICs |
| N-Acetyl-<br>aspartic acid | Uridine<br>monophosphate | MassIVE/<br>MSV000086399 | *C. elegans* | RPC | - |  | Detailed EICs |
| Arabitol | Dihydroxypr<br>opanoic<br>acid | MetaboLights/<br>MTBLS464 | *E. coli* | HILIC | [29] |  | Detailed EICs |
| Proline | Threoninyl-Aspartate | MassIVE/<br>MSV000083073 | *R.*<br>*norvegicus* | RPC | - |  | Detailed EICs |
| Proline | Threoninyl-Aspartate | Metabolomics/<br>Workbench<br>ST001514 | Phyto-<br>plankton | HILIC | [30] |  | Detailed EICs |

Moreover, it is worth reporting that we also conducted a MASST search of MS2 spectra generated for the heteroadducts detected in the yeast data, which attempts to find these MS2 spectra within the raw data deposited on MassIVE. However, this search did not return any results, which supports our previous statement, that heteromers rarely trigger MS2 scans due to their low abundance, with state-of-the-art MS2-acquisition strategies.

#### 3.6.3 Data availability and deposition of MS2 scans in GNPS library

To help other researchers in avoiding misidentification of multimeric adducts in their own datasets, we submitted the acquired MS2 spectra for annotated multimeric adducts for which we were able to annotate the associated monomers in the GNPS public MS2 library. Moreover, raw data (MS1 and MS2) were released to MassIVE (Dataset ID: MSV000089018)

## 4. Conclusions

We have developed a workflow to recognize multimeric molecular species in metabolomics data. While the implementation of the presented method is limited by the availability of fully 13C labeled organisms several important findings are of interest for non-targeted metabolomics studies in general. First, we found that for four different data sets of different complexity, the number of heteroadducts did not exceed 5% of all detectable biological features. We also observed results suggesting that HILIC is more prone to heteromer formation than RPC. Moreover, we showed that heteroadducts are generally observed to be less abundant than the median of metabolite peaks. Still, continuous progress in MS2-acquisition coverage will likely increase the frequency in which multimer-MS2 spectra are observed in metabolomics data.

This is supported by the fact that we were able to find convincing evidence that heteroadducts identified in this study are also present in public datasets produced by other labs from the analysis of other biological extracts. As we do not expect our workflow to be routinely employable due to the limited availability of fully 13C labeled sample material, we compiled an MS2 library of the detected heteroadducts. This library can be readily used in future non-targeted metabolomics studies by other groups.

## Acknowledgment

The authors would like to thank Gerrit Hermann from ISOtopic Solutions for providing the labeled and unlabeled yeast material.

## References

[1] R.R. da Silva, P.C. Dorrestein, R.A. Quinn, Illuminating the dark matter in metabolomics, Proc. Natl. Acad. Sci. 112 (2015) 12549–12550. https://doi.org/10.1073/pnas.1516878112.

[2] C. Kuhl, R. Tautenhahn, C. Böttcher, T.R. Larson, S. Neumann, CAMERA: An Integrated Strategy for Compound Spectra Extraction and Annotation of Liquid Chromatography/Mass Spectrometry Data Sets, Anal. Chem. 84 (2012) 283–289. https://doi.org/10.1021/ac202450g.

[3] C.D. Broeckling, F.A. Afsar, S. Neumann, A. Ben-Hur, J.E. Prenni, RAMClust: A Novel Feature Clustering Method Enables Spectral-Matching-Based Annotation for Metabolomics Data, Anal. Chem. 86 (2014) 6812–6817. https://doi.org/10.1021/ac501530d.

[4] R. Schmid, D. Petras, L.-F. Nothias, M. Wang, A.T. Aron, A. Jagels, H. Tsugawa, J. Rainer, M. Garcia-Aloy, K. Dührkop, A. Korf, T. Pluskal, Z. Kameník, A.K. Jarmusch, A.M. Caraballo-Rodríguez, K.C. Weldon, M. Nothias-Esposito, A.A. Aksenov, A. Bauermeister, A. Albarracin Orio, C.O. Grundmann, F. Vargas, I. Koester, J.M. Gauglitz, E.C. Gentry, Y. Hövelmann, S.A. Kalinina, M.A. Pendergraft, M. Panitchpakdi, R. Tehan, A. Le Gouellec, G. Aleti, H. Mannochio Russo, B. Arndt, F. Hübner, H. Hayen, H. Zhi, M. Raffatellu, K.A. Prather, L.I. Aluwihare, S. Böcker, K.L. McPhail, H.-U. Humpf, U. Karst, P.C. Dorrestein, Ion identity molecular networking for mass spectrometry-based metabolomics in the GNPS environment, Nat. Commun. 12 (2021) 3832. https://doi.org/10.1038/s41467-021-23953-9.

[5] N.G. Mahieu, J.L. Spalding, S.J. Gelman, G.J. Patti, Defining and Detecting Complex Peak Relationships in Mass Spectral Data: The Mz.unity Algorithm, Anal. Chem. 88 (2016) 9037–9046. https://doi.org/10.1021/acs.analchem.6b01702.

[6] N.G. Mahieu, X. Huang, Y.-J. Chen, G.J. Patti, Credentialing Features: A Platform to Benchmark and Optimize Untargeted Metabolomic Methods, Anal. Chem. 86 (2014) 9583–9589. https://doi.org/10.1021/ac503092d.

[7] N.G. Mahieu, G.J. Patti, Systems-Level Annotation of a Metabolomics Data Set Reduces 25 000 Features to Fewer than 1000 Unique Metabolites, Anal. Chem. 89 (2017) 10397–10406. https://doi.org/10.1021/acs.analchem.7b02380.

[8] K. Ortmayr, M. Schwaiger, S. Hann, G. Koellensperger, An integrated metabolomics workflow for the quantification of sulfur pathway intermediates employing thiol protection with N-ethyl maleimide and hydrophilic interaction liquid chromatography tandem mass spectrometry, Analyst. 140 (2015) 7687–7695. https://doi.org/10.1039/C5AN01629K.

[9] C. Haberhauer-Troyer, M. Delic, B. Gasser, D. Mattanovich, S. Hann, G. Koellensperger, Accurate quantification of the redox-sensitive GSH/GSSG ratios in the yeast Pichia pastoris by HILIC–MS/MS, Anal. Bioanal. Chem. 405 (2013) 2031–2039. https://doi.org/10.1007/s00216-012-6620-4.

[10] C. Sauerschnig, M. Doppler, C. Bueschl, R. Schuhmacher, Methanol Generates Numerous Artifacts during Sample Extraction and Storage of Extracts in Metabolomics Research, Metabolites. 8 (2018) 1. https://doi.org/10.3390/metabo8010001.

[11] S. Neubauer, C. Haberhauer-Troyer, K. Klavins, H. Russmayer, M.G. Steiger, B. Gasser, M. Sauer, D. Mattanovich, S. Hann, G. Koellensperger, U13C cell extract of Pichia pastoris – a powerful tool for evaluation of sample preparation in metabolomics, J. Sep. Sci. 35 (2012) 3091–3105. https://doi.org/10.1002/jssc.201200447.

[12] M. Schwaiger, H. Schoeny, Y.E. Abiead, G. Hermann, E. Rampler, G. Koellensperger, Merging metabolomics and lipidomics into one analytical run, Analyst. 144 (2018) 220–229. https://doi.org/10.1039/C8AN01219A.

[13] C. Bueschl, B. Kluger, N.K.N. Neumann, M. Doppler, V. Maschietto, G.G. Thallinger, J. Meng-Reiterer, R. Krska, R. Schuhmacher, MetExtract II: A Software Suite for Stable Isotope-Assisted Untargeted Metabolomics, Anal. Chem. 89 (2017) 9518–9526. https://doi.org/10.1021/acs.analchem.7b02518.

[14] M.C. Chambers, B. Maclean, R. Burke, D. Amodei, D.L. Ruderman, S. Neumann, L. Gatto, B. Fischer, B. Pratt, J. Egertson, K. Hoff, D. Kessner, N. Tasman, N. Shulman, B. Frewen, T.A. Baker, M.-Y. Brusniak, C. Paulse, D. Creasy, L. Flashner, K. Kani, C. Moulding, S.L. Seymour, L.M. Nuwaysir, B. Lefebvre, F. Kuhlmann, J. Roark, P. Rainer, S. Detlev, T. Hemenway, A. Huhmer, J. Langridge, B. Connolly, T. Chadick, K. Holly, J. Eckels, E.W. Deutsch, R.L. Moritz, J.E. Katz, D.B. Agus, M. MacCoss, D.L. Tabb, P. Mallick, A cross-platform toolkit for mass spectrometry and proteomics, Nat. Biotechnol. 30 (2012) 918–920. https://doi.org/10.1038/nbt.2377.

[15] B. Seidl, R. Schuhmacher, C. Bueschl, CPExtract, a Software Tool for the Automated Tracer-Based Pathway Specific Screening of Secondary Metabolites in LC-HRMS Data, Anal. Chem. 94 (2022) 3543–3552. https://doi.org/10.1021/acs.analchem.1c04530.

[16] M. Wang, J.J. Carver, V.V. Phelan, L.M. Sanchez, N. Garg, Y. Peng, D.D. Nguyen, J. Watrous, C.A. Kapono, T. Luzzatto-Knaan, C. Porto, A. Bouslimani, A.V. Melnik, M.J. Meehan, W.-T. Liu, M. Crüsemann, P.D. Boudreau, E. Esquenazi, M. Sandoval-Calderón, R.D. Kersten, L.A. Pace, R.A. Quinn, K.R. Duncan, C.-C. Hsu, D.J. Floros, R.G. Gavilan, K. Kleigrewe, T. Northen, R.J. Dutton, D. Parrot, E.E. Carlson, B. Aigle, C.F. Michelsen, Jelsbak, C. Sohlenkamp, P. Pevzner, A. Edlund, J. McLean, J. Piel, B.T. Murphy, L. Gerwick, C.-C. Liaw, Y.-L. Yang, H.-U. Humpf, M. Maansson, R.A. Keyzers, A.C. Sims, A.R. Johnson, A.M. Sidebottom, B.E. Sedio, A. Klitgaard, C.B. Larson, C.A. Boya P, D. Torres-Mendoza, D.J. Gonzalez, D.B. Silva, L.M. Marques, D.P. Demarque, E. Pociute, E.C. O’Neill, E. Briand, E.J.N. Helfrich, E.A. Granatosky, E. Glukhov, F. Ryffel, H. Houson, H. Mohimani, J.J. Kharbush, Y. Zeng, J.A. Vorholt, K.L. Kurita, P. Charusanti, K.L. McPhail, K.F. Nielsen, L. Vuong, M. Elfeki, M.F. Traxler, N. Engene, N. Koyama, O.B. Vining, R. Baric, R.R. Silva, S.J. Mascuch, S. Tomasi, S. Jenkins, V. Macherla, T. Hoffman, V. Agarwal, P.G. Williams, J. Dai, R. Neupane, J. Gurr, A.M.C. Rodríguez, A. Lamsa, C. Zhang, K. Dorrestein, B.M. Duggan, J. Almaliti, P.-M. Allard, P. Phapale, L.-F. Nothias, T. Alexandrov, M. Litaudon, J.-L. Wolfender, J.E. Kyle, T.O. Metz, T. Peryea, D.-T. Nguyen, D. VanLeer, P. Shinn, A. Jadhav, R. Müller, K.M. Waters, W. Shi, X. Liu, L. Zhang, R. Knight, P.R. Jensen, B.Ø. Palsson, K. Pogliano, R.G. Linington, M. Gutiérrez, N.P. Lopes, W.H. Gerwick, B.S. Moore, P.C. Dorrestein, N. Bandeira, Sharing and community curation of mass spectrometry data with Global Natural Products Social Molecular Networking, Nat. Biotechnol. 34 (2016) 828–837. https://doi.org/10.1038/nbt.3597.

[17] K. Dührkop, M. Fleischauer, M. Ludwig, A.A. Aksenov, A.V. Melnik, M. Meusel, P.C. Dorrestein, J. Rousu, S. Böcker, SIRIUS 4: a rapid tool for turning tandem mass spectra into metabolite structure information, Nat. Methods. 16 (2019) 299–302. https://doi.org/10.1038/s41592-019-0344-8.

[18] F. Wang, J. Liigand, S. Tian, D. Arndt, R. Greiner, D.S. Wishart, CFM-ID 4.0: More Accurate ESI-MS/MS Spectral Prediction and Compound Identification, Anal. Chem. 93 (2021) 11692–11700. https://doi.org/10.1021/acs.analchem.1c01465.

[19] A. Ćeranić, M. Doppler, C. Büschl, A. Parich, K. Xu, A. Koutnik, H. Bürstmayr, M. Lemmens, R. Schuhmacher, Preparation of uniformly labelled 13C- and 15N-plants using customised growth chambers, Plant Methods. 16 (2020) 46. https://doi.org/10.1186/s13007-020-00590-9.

[20] B. Warth, A. Parich, C. Bueschl, D. Schoefbeck, N.K.N. Neumann, B. Kluger, K. Schuster, R. Krska, G. Adam, M. Lemmens, R. Schuhmacher, GC–MS based targeted metabolic profiling identifies changes in the wheat metabolome following deoxynivalenol treatment, Metabolomics. 11 (2015) 722–738. https://doi.org/10.1007/s11306-014-0731-1.

[21] M. Wang, A.K. Jarmusch, F. Vargas, A.A. Aksenov, J.M. Gauglitz, K. Weldon, D. Petras, R. da Silva, R. Quinn, A.V. Melnik, J.J.J. van der Hooft, A.M. Caraballo-Rodríguez, L.F. Nothias, C.M. Aceves, M. Panitchpakdi, E. Brown, F. Di Ottavio, N. Sikora, E.O. Elijah, L. Labarta-Bajo, E.C. Gentry, S. Shalapour, K.E. Kyle, S.P. Puckett, J.D. Watrous, C.S. Carpenter, A. Bouslimani, M. Ernst, A.D. Swafford, E.I. Zúñiga, M.J. Balunas, J.L. Klassen, R. Loomba, R. Knight, N. Bandeira, P.C. Dorrestein, Mass spectrometry searches using MASST, Nat. Biotechnol. 38 (2020) 23–26. https://doi.org/10.1038/s41587-019-0375-9.

[22] K. Haug, K. Cochrane, V.C. Nainala, M. Williams, J. Chang, K.V. Jayaseelan, C. O’Donovan, MetaboLights: a resource evolving in response to the needs of its scientific community, Nucleic Acids Res. 48 (2020) D440–D444. https://doi.org/10.1093/nar/gkz1019.

[23] M. Sud, E. Fahy, D. Cotter, K. Azam, I. Vadivelu, C. Burant, A. Edison, O. Fiehn, R. Higashi, K.S. Nair, S. Sumner, S. Subramaniam, Metabolomics Workbench: An international repository for metabolomics data and metadata, metabolite standards, protocols, tutorials and training, and analysis tools, Nucleic Acids Res. 44 (2016) D463–D470. https://doi.org/10.1093/nar/gkv1042.

[24] D. Petras, V.V. Phelan, D. Acharya, A.E. Allen, A.T. Aron, N. Bandeira, B.P. Bowen, D. Belle-Oudry, S. Boecker, D.A. Cummings, J.M. Deutsch, E. Fahy, N. Garg, R. Gregor, J. Handelsman, M. Navarro-Hoyos, A.K. Jarmusch, S.A. Jarmusch, K. Louie, K.N. Maloney, M.T. Marty, M.M. Meijler, I. Mizrahi, R.L. Neve, T.R. Northen, C. Molina-Santiago, M. Panitchpakdi, B. Pullman, A.W. Puri, R. Schmid, S. Subramaniam, M. Thukral, F. Vasquez-Castro, P.C. Dorrestein, M. Wang, GNPS Dashboard: collaborative exploration of mass spectrometry data in the web browser, Nat. Methods. 19 (2022) 134–136. https://doi.org/10.1038/s41592-021-01339-5.

[25] R. Giné, J. Capellades, J.M. Badia, D. Vughs, M. Schwaiger-Haber, T. Alexandrov, M. Vinaixa, A.M. Brunner, G.J. Patti, O. Yanes, HERMES: a molecular-formula-oriented method to target the metabolome, Nat. Methods. 18 (2021) 1370–1376. https://doi.org/10.1038/s41592-021-01307-z.

[26] Z. Zuo, L. Cao, L.-F. Nothia, H. Mohimani, MS2Planner: improved fragmentation spectra coverage in untargeted mass spectrometry by iterative optimized data acquisition, Bioinformatics. 37 (2021) i231–i236. https://doi.org/10.1093/bioinformatics/btab279.

[27] M. Yu, G. Dolios, L. Petrick, Reproducible untargeted metabolomics workflow for exhaustive MS2 data acquisition of MS1 features, J. Cheminformatics. 14 (2022) 6. https://doi.org/10.1186/s13321-022-00586-8.

[28] A.K. Jarmusch, M. Wang, C.M. Aceves, R.S. Advani, S. Aguirre, A.A. Aksenov, G. Aleti, A.T. Aron, A. Bauermeister, S. Bolleddu, A. Bouslimani, A.M. Caraballo Rodriguez, R. Chaar, R. Coras, E.O. Elijah, M. Ernst, J.M. Gauglitz, E.C. Gentry, M. Husband, S.A. Jarmusch, K.L. Jones, Z. Kamenik, A. Le Gouellec, A. Lu, L.-I. McCall, K.L. McPhail, M.J. Meehan, A.V. Melnik, R.C. Menezes, Y.A. Montoya Giraldo, N.H. Nguyen, L.F. Nothias, M. Nothias-Esposito, M. Panitchpakdi, D. Petras, R.A. Quinn, N. Sikora, J.J.J. van der Hooft, F. Vargas, A. Vrbanac, K.C. Weldon, R. Knight, N. Bandeira, P.C. Dorrestein, ReDU: a framework to find and reanalyze public mass spectrometry data, Nat. Methods. 17 (2020) 901–904. https://doi.org/10.1038/s41592-020-0916-7.

[29] I.M. Vincent, D.E. Ehmann, S.D. Mills, M. Perros, M.P. Barrett, Untargeted Metabolomics To Ascertain Antibiotic Modes of Action, Antimicrob. Agents Chemother. 60 (n.d.) 2281–2291. https://doi.org/10.1128/AAC.02109-15.

[30] K.R. Heal, B.P. Durham, A.K. Boysen, L.T. Carlson, W. Qin, F. Ribalet, A.E. White, R.M. Bundy, E.V. Armbrust, A.E. Ingalls, Marine Community Metabolomes Carry Fingerprints of Phytoplankton Community Composition, MSystems. 6 (n.d.) e01334–20. https://doi.org/10.1128/mSystems.01334-20.

