## Supplementary Material 1-4_AND_Figure S1 for "Heterogeneous multimeric metabolite ion species observed in LC-MS based metabolomics data sets"

### Supplementary Information 1

##### Wheat ear dataset

Wheat ear samples were produced as described in (Ćeranić et al. 2020).
Briefly summarized: Wheat plants of the genotype "Remus" were grown in a hydroponic setup in hermetically sealed growth chambers in parallel. The conditions in both boxes were kept identically, however, in one box the atmosphere consisted of native $CO_2$ (for the generation of the native or unlabeled wheat plants), while in the other box the atmosphere consisted of ~99% $^{13}CO_2$. Wheat plants were treated with water (mock) at flowering stage and samples were harvested 96 hours after treatment (Warth et al. 2015). For this, the treated spikelets and the rachis were cut from the wheat ear, put in liquid nitrogen, and subsequently frozen at -80°C until further analysis.

Samples were prepared for LC-HRMS analysis as described in (Ćeranić et al. 2020). In short, wheat ears were ground into a fine powder and 100 mg of the frozen powder was extracted in 1 ml MeOH/$H_2O$ (3:1, v/v) + 0.1% FA. The samples were vortexed, put into an ultrasonic bath for 15 min, and vortexed again for 10 min at 14,000 rpm. The supernatants were diluted with $H_2O$ + 0.1% FA to achieve a ratio of MeOH/$H_2O$ (1:1, v/v) + 0.1% FA.  At this stage the extracts of the native and $^{13}C$-labeled wheat plant extracts were mixed in a 1:1 ratio for LC-HRMS analysis.

### Supplementary Information 2

##### *Fusarium graminearum* cultivation

*Fusarium graminearum* PH-I (wildtype) and two *F.g.* Δkmt6  strains were grown on liquid minimal medium (1.0 g·L$^{-1}$ $KH_2HPO_4$, 0.5 g·L$^{-1}$ $MgSO_4 \cdot 7 H_2O$, 0.5 g·L$^{-1}$ KCl, 0.48 g·L$^{-1}$ $NH_4NO_3$ and 0.2mL·L$^{-1}$ Vogel's trace solution as described in (Vogel, 1956)) containing 1 % (w·v$^{-1}$) D-glucose (native glucose) or  U-$^{13}C_6$ labeled D-glucose in 10 mL GC headspace glass vials in a volume of 2.0 mL each. In each case 2 mL medium was inoculated with 12 µL fresh spore solution (final concentration 25,000 spores·mL$^{-1}$). The vials were sealed with a cellulose stopper during incubation. Cultures were incubated in complete darkness at 18 °C and 70% relative humidity for 14 and 21 days. Three biological replicates were made for each approach.

##### *Trichoderma reesei* cultivation

*Trichoderma reesei*  (strains QM6a (WT), ΔXpp1, ΔXpp2 and ΔXpp1ΔXpp2) were grown on a liquid minimal medium based on Mandels-Andreotti (MA) medium as described in (Mandels, 1985) without peptone containing 1 % (w·v$^{-1}$) D-glucose or U-$^{13}C_6$ labeled glucose as the sole carbon source in 24 well cell culture plates in a volume of 1.0 mL per well. In each case 1 mL medium was  inoculated with 100 µL spore solution (0.05 absorption units at $OD_{700}$ at 1 cm path length) at 30 °C and 80 % relative humidity in complete darkness for 96 hours. Six  biological replicates were made for each approach.

##### Sample preparation

The fungal mycelium was removed using a pipette tip and the culture supernatant was filtered through glass wool in each case to remove mycelium leftovers. The filtered supernatant was cooled on ice and quenched by adding 30 % (v·v$^{-1}$) cold (-20 °C) acetonitrile, centrifuged at 30,000 x g for 20 minutes at 4 °C and transferred into HPLC vials for LC-HRMS analysis.

### Supplementary Information 3

**LC-HRMS analysis**

Mixture samples of native and $^{13}C$ labeled material were analyzed with an QExactive HF Orbitrap instrument coupled to a Vanquish UHPLC system (Thermo Fisher Scientific). The analysis is detailed in (Sauerschnig et al. 2018).
Briefly summarized: An X-Bridge $C_{18}$-column (150 x 2.1 mm i.d., 3.5 µm particle size – Waters, Milford, MA, USA) was employed for reversed-phase chromatographic separation. H2O + 0.1% FA (eluent A) and MeOH + 0.1% FA (eluent B) were used in a linear gradient (0 min: 10% B; 2 min: 10% B; 32 min: 100% B; 37 min: 100% B; 45 min: 10% B) lasting for 45 minutes with a constant flow rate of 250 µl $min^{-1}$. Ionization was carried out with a heated electrospray ionization (HESI) source and fast polarity switching was employed (ionization was switched after each scan). MS1 data was acquired with a resolution of 120,000 (FWHM at *m/z* 200).

### Supplementary Information 4

**Data processing with MetExtract II and CPExtract**

Raw LC-HRMS data was converted to the mzXML format with ProteoWizard. Then, the same data processing was applied as for the yeast dataset detailed in the main manuscript.
The following parameter values were chosen differently than for the yeast dataset for MetExtract II and CPExtract data evaluation:

<u>General parameters</u>

Cn count to search for: 3 – 60, retention time domain to search for: 3 – 36 minutes (RP only method), minimum intensity threshold for signals: 1E4 (only applicable for the monoisotopic and the fully $^{13}C$ labeled principal isotopologues), maximum allowed mass deviation for isotopologues in the same scan: 3 ppm, isotopologue peaks checked: M + 1 and M + Cn-1, maximum cluster ppm for region of interest (ROI) generation: 8 ppm, minimum number of signals with verified isotopologue pattern: 3, chromatographic peak width in wavelet terms: 3 - 19, bracketing mz and retention-time window: 10 ppm and 0.1 minutes, minimum Pearson correlation coefficient (PCC) for feature grouping were 0.85

<u>Specific parameters of the Wheat dataset</u>

$^{13}C$-isotopic enrichment: 98.7%, minimum intensity threshold for signals: 1E6 (only applicable for the monoisotopic and the fully $^{13}C$ labeled principal isotopologues)

<u>Specific parameters of the *Fusarium graminearum* dataset</u>

$^{13}C$-isotopic enrichment: 98.4%, minimum intensity threshold for signals: 1E4 (only applicable for the monoisotopic and the fully $^{13}C$ labeled principal isotopologues)

<u>Specific parameters of the *Trichoderma reesei* dataset</u>

$^{13}C$-isotopic enrichment: 98.9%, minimum intensity threshold for signals: 1E4 (only applicable for the monoisotopic and the fully $^{13}C$ labeled principal isotopologues)

### References


Ćeranić, A., Doppler, M., Büschl, C., Parich, A., Xu, K., Koutnik, A., Bürstmayr, H., Lemmens, M., Schuhmacher, R.. 2020. “Preparation of Uniformly Labelled 13C- and 15N-Plants Using Customised Growth Chambers.” Plant Methods 16 (1): 46. https://doi.org/10.1186/s13007-020-00590-9

Sauerschnig, C., Doppler, M., Bueschl, C., Schuhmacher, R.. 2018. “Methanol Generates Numerous Artifacts during Sample Extraction and Storage of Extracts in Metabolomics Research.” *Metabolites* 8 (1): 1. https://doi.org/10.3390/metabo8010001

Warth, B., Parich, A., Bueschl, C., Schoefbeck, D., Neumann, NKN., Kluger, B., Schuster, K., et al. 2015. “GC–MS Based Targeted Metabolic Profiling Identifies Changes in the Wheat Metabolome Following Deoxynivalenol Treatment.” *Metabolomics* 11 (3): 722–38. https://doi.org/10.1007/s11306-014-0731-1

Vogel, HJ.. 1956. “A convenient growth medium for Neurospora (medium N)” *Microb. Genet. Bull.* 13, 42-43.

Mandels, M.. 1985. “Applications of cellulases” *Biochemical Society Transactions* 13 (2), 414-416.


### Supplementary Figures

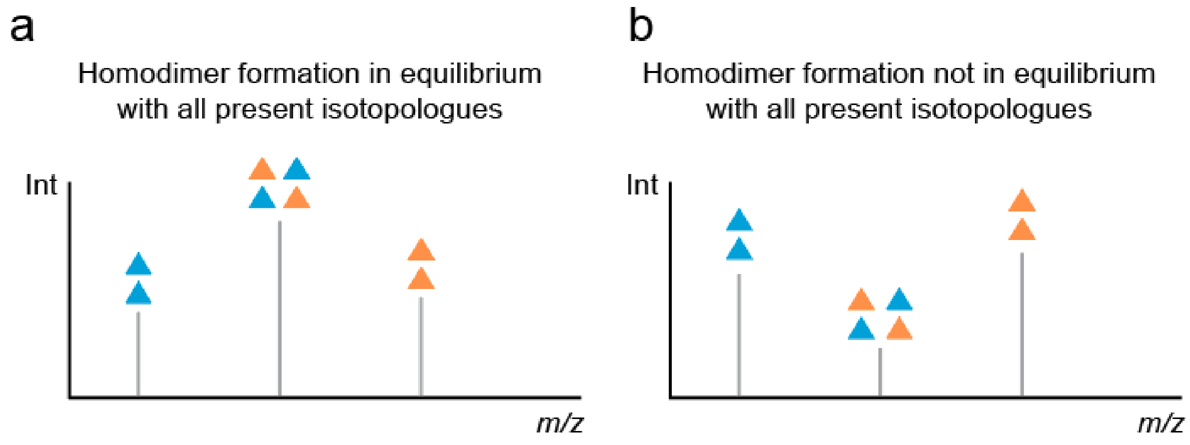


**Figure S1 Isotopologue pattern observed for homodimers.** (a) Dimer formed when 13C and 12C extracts were already mixed. (b) Dimer formed partly before 13C and 12C extracts were mixed.
